# *Xenopus* hybrids provide insight into cell and organism size control

**DOI:** 10.1101/395814

**Authors:** Romain Gibeaux, Kelly Miller, Rachael Acker, Taejoon Kwon, Rebecca Heald

## Abstract

Determining how size is controlled is a fundamental question in biology that is poorly understood at the organismal, cellular and subcellular levels. The *Xenopus* species, *X. laevis* and *X. tropicalis* differ in size at all three of these levels. Despite these differences, fertilization of *X. laevis* eggs with *X. tropicalis* sperm gives rise to viable hybrid animals that are intermediate in size. We observed that although hybrid and *X. laevis* embryogenesis initiates from the same sized zygote and proceeds synchronously through development, hybrid animals were smaller by the tailbud stage, and a change in the ratio of nuclear size to cell size was observed shortly after zygotic genome activation (ZGA), suggesting that differential gene expression contributes to size differences. Transcriptome analysis at the onset of ZGA identified twelve transcription factors paternally expressed in hybrids. A screen of these *X. tropicalis* factors by expression in *X. laevis* embryos revealed that Hes7 and Ventx2 significantly reduced *X. laevis* body length size by the tailbud stage, although nuclear to cell size scaling relationships were not affected as in the hybrid. Together, these results suggest that transcriptional regulation contributes to biological size control in *Xenopus*.

## Introduction

Biological size control and scaling are important and fundamental features of living systems. However, the molecular mechanisms that control size the organism, cell, and subcellular level are poorly understood. The frog *Xenopus* has emerged as a powerful system to explore nuclear and spindle size differences that occur between related species with different-sized eggs (Kitaoka et al., 2018; Levy and Heald, 2010; Loughlin et al., 2011), as well as subcellular scaling during early development, when cleavage divisions cause a rapid reduction in cell size (Good et al., 2013; Wilbur and Heald, 2013). We therefore set out to investigate whether *Xenopus* frogs could also be used to study size control at the level of the cell and the whole organism.

Cell size correlates strongly and linearly with genome size in a myriad of different organisms (Cavalier-Smith, 2005; Gregory, 2001; Mirsky, 1951), and increases in genome copy number through polyploidy have been shown to increase cell size within tissues or cell types (Frawley and Orr-Weaver, 2015; Lee et al., 2009). However, the molecular link between genome size and cell size remains an open question. Although increases in ploidy may globally affect gene expression, work in unicellular organisms such as yeast suggests that the maintenance of scaling between genome size and cell size does not simply reflect gene dosage (Galitski et al., 1999; Marguerat et al., 2012; Neumann and Nurse, 2007). Furthermore, the correlation between genome size and cell size is independent of the proportion of the genome that codes for genes (Cavalier-Smith, 2005; Gregory, 2001; Taft et al., 2007). A number of factors involved in many different processes, such as growth, metabolism and protein synthesis, development, differentiation, and cell cycle regulation (Björklund et al., 2006) can influence cell size in a variety of organisms, from bacteria, to yeast, to *Drosophila*, to mammals (Marguerat and Bähler, 2012). Many of these genes are conserved and contribute to tissue and organ size in a variety of multicellular organisms, however, how they influence organism size, and how organism size feeds back to organ/tissue/cell size to attain homeostasis remains unclear.

Interestingly, in the related frog species *Xenopus laevis* and *Xenopus tropicalis*, the size of the genome, cells, and component subcellular structures scale with body size (Brown et al., 2007; Levy and Heald, 2010). Furthermore, the larger allotetraploid *Xenopus laevis* (6.2 × 10^9^ base pairs, N = 36 chromosomes, average body length 10 cm) and smaller diploid *Xenopus tropicalis* (3.4 x 10^9^ base pairs, N = 20 chromosomes, 4 cm in length) can hybridize. While fertilization of an *X. tropicalis* egg with a *X. laevis* sperm produces an inviable hybrid embryo that dies as a late blastula (Gibeaux et al., 2018), fertilization of an *X. laevis* egg with a *X. tropicalis* sperm (*l*_e_×*t*_s_) produces a viable adult frog intermediate in genome size (N = 28 chromosomes) and body length between the two species (Narbonne et al., 2011). This viable hybrid thus provides a unique *in vivo* vertebrate model for investigating biological size control at the organismal, cellular, and subcellular levels.

In this study, we characterized size scaling in viable *l*_e_×*t*_s_ hybrids and used this system to establish a novel screening method for candidate genes involved in size control to identify factors that affect the body size of the frog, as well as the scaling of its component cells and subcellular structures.

## Results

### Reduced size in viable *l*_e_×*t*_s_ hybrids

Whereas cross-fertilization of *X. tropicalis* eggs with *X. laevis* sperm produces hybrid embryos that die during zygotic genome activation (ZGA), the reverse cross of *X. laevis* eggs and *X. tropicalis* sperm (*l*_e_×*t*_s_) results in viable hybrid embryos that possess genetic features of both *X. laevis* and *X. tropicalis* parents (Bürki, 1985; Elurbe et al., 2017; Gibeaux et al., 2018; Lindsay et al., 2003; Narbonne et al., 2011). Hybrid embryos progressed through tadpole, froglet, and adult stages (Figure 1A, B), although with significant morbidity. Using standard husbandry conditions for *X. laevis*, only four adults were obtained from hundreds of embryos in two separate attempts to generate *l*_e_×*t*_s_ hybrid frogs. Early development in the *l*_e_×*t*_s_ hybrid proceeded normally according to Nieuwkoop and Faber staging (Nieuwkoop and Faber, 1994) until the end of neurulation, and at a similar rate compared to wild type *X. laevis* embryos (Figure 1C, Supplementary Movie S1). However, by the tailbud stage, body length was significantly decreased in *l*_e_×*t*_s_ hybrids (Figure 1D). Relative shortening of body length continued, although development remained similar to *X. laevis*, and both hybrid and control animals initiated metamorphosis with the same timing (Figure 1E). As soon as metamorphosis was complete, size scaling stopped and the body length of both the *l*_e_×*t*_s_ hybrid and *X. laevis* froglets increased at the same rate, retaining the difference in size (Figure 1F). Strikingly, in adult hybrid frogs, both cell and nuclear size of erythrocytes was reduced (Figure 1G). Since the hybrid genome (28 chromosomes) is smaller than the *X. laevis* genome (36 chromosomes), these observations are consistent with genome size-dependent scaling at the organism, cellular and subcellular levels.

**Figure 1.**
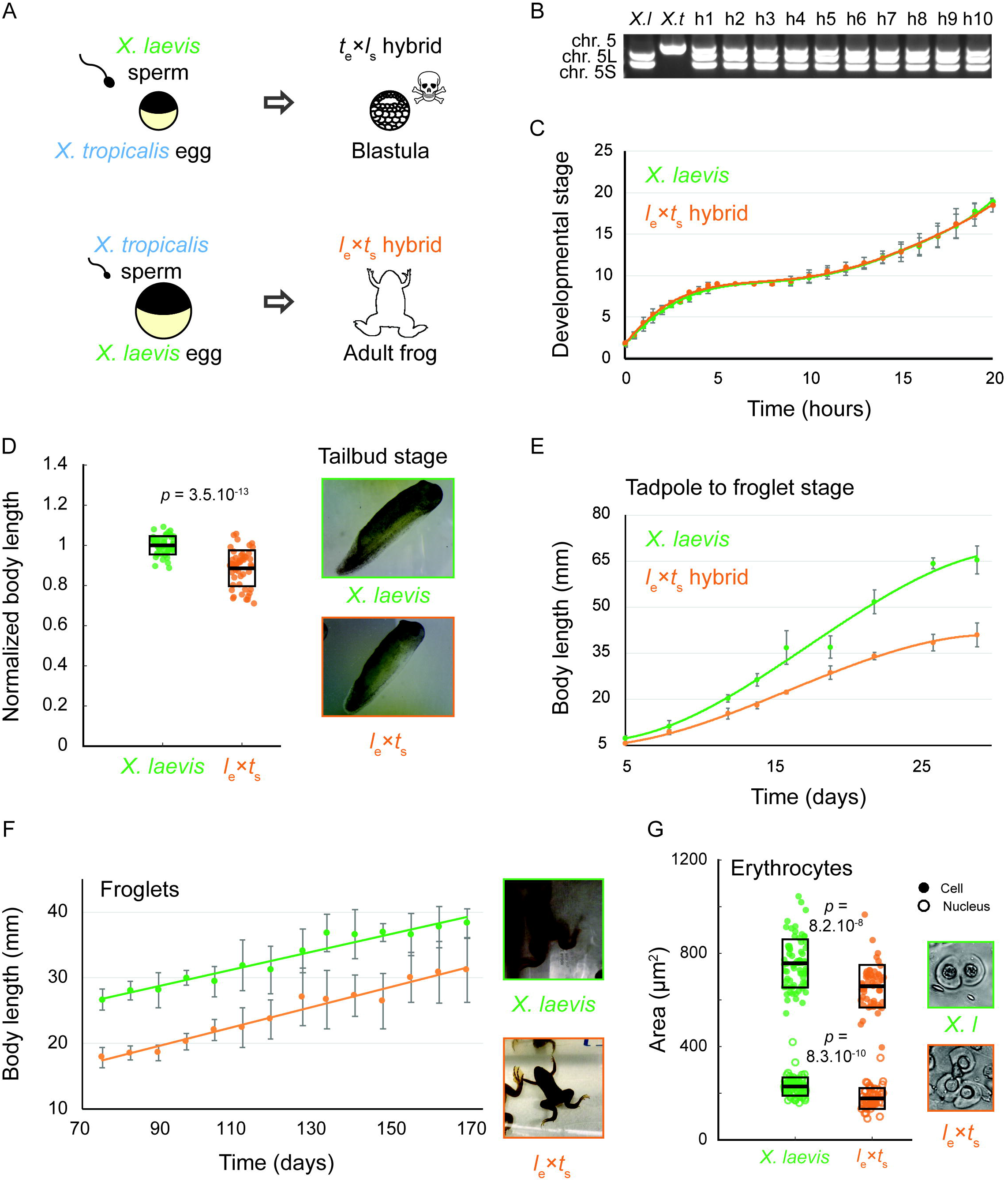
Growth and development of *Xenopus l*_e_×*t*_s_ viable hybrids. **(A)** Schematic of developmental outcomes of *Xenopus laevis* and *Xenopus tropicalis* cross-fertilization. **(B)** Agarose gel electrophoresis showing PCR amplification of 2 genomic loci in *X. laevis* (*X. l*, on chromosomes 5L and 5S) and one locus in *X. tropicalis* (*X. t*, chromosome 5). H1-10 indicates 10 randomly chosen hybrid tadpoles tested, confirming the consistent presence of all 3 subgenomes in hybrids. **(C)** Developmental timing in *X. laevis* and *l*_e_×*t*_s_ hybrid embryos. Average is plotted for each time point. Error bars show standard deviation. **(D)** Body length of tailbud stage *X. laevis* and *l*_e_×*t*_s_ hybrids. Box plots show all individual body lengths. Thick line inside box = average length, upper and lower box boundaries = +/- standard deviation (SD). P-value was determined by two-tailed heteroscedastic t-test. Representative images of tailbuds at identical scale are shown on the right. **(E)** Body length of tadpoles throughout metamorphosis for *X. laevis* and *l*_e_×*t*_s_ hybrids. Average is plotted for each time point. Error bars show standard deviation. **(F)** Body length of *X. laevis* and *l*_e_×*t*_s_ hybrid froglets. Average is plotted for each time point. Error bars show standard deviation. Representative images of froglets at identical scale are shown on the right. **(G)** Size of erythrocyte cells and nuclei in *X. laevis* and *l*_e_×*t*_s_ hybrid adult frogs. Box plots show all individual cell or nuclear areas. Thick line inside box = average area, upper and lower box boundaries = +/- SD. P-values were determined by two-tailed heteroscedastic t-test. Representative images of erythrocytes at identical scale are shown on the right.

### Nuclear to cell size scaling in *l*_e_×*t*_s_ hybrids is more similar to that of *X. laevis* haploids, despite a larger genome size

We wondered whether the scaling observed in hybrid embryos was due to the decrease in genome size, or if the paternal *X. tropicalis* genome also influenced size scaling. To examine the effect of altering genome size alone, we utilized haploid *X. laevis* embryos produced by fertilizing wild type *X. laevis* eggs with irradiated *X. laevis* sperm. While the sperm DNA is inactivated and does not contribute to the genome of the offspring, the sperm centrosome induces embryonic development yielding haploid embryos containing only the N=18 maternal genome (Hamilton, 1957). Haploid embryos developed normally to the tailbud stage, and at a similar developmental rate to wild type *X. laevis* embryos (Figure 2B, Supplementary Movie S2). However, by the tailbud stage, body length was significantly reduced in *X. laevis* haploids (Figure 2C). Haploid embryos never reach metamorphosis and stop developing as stunted tadpoles (Hamilton, 1963).

**Figure 2.**
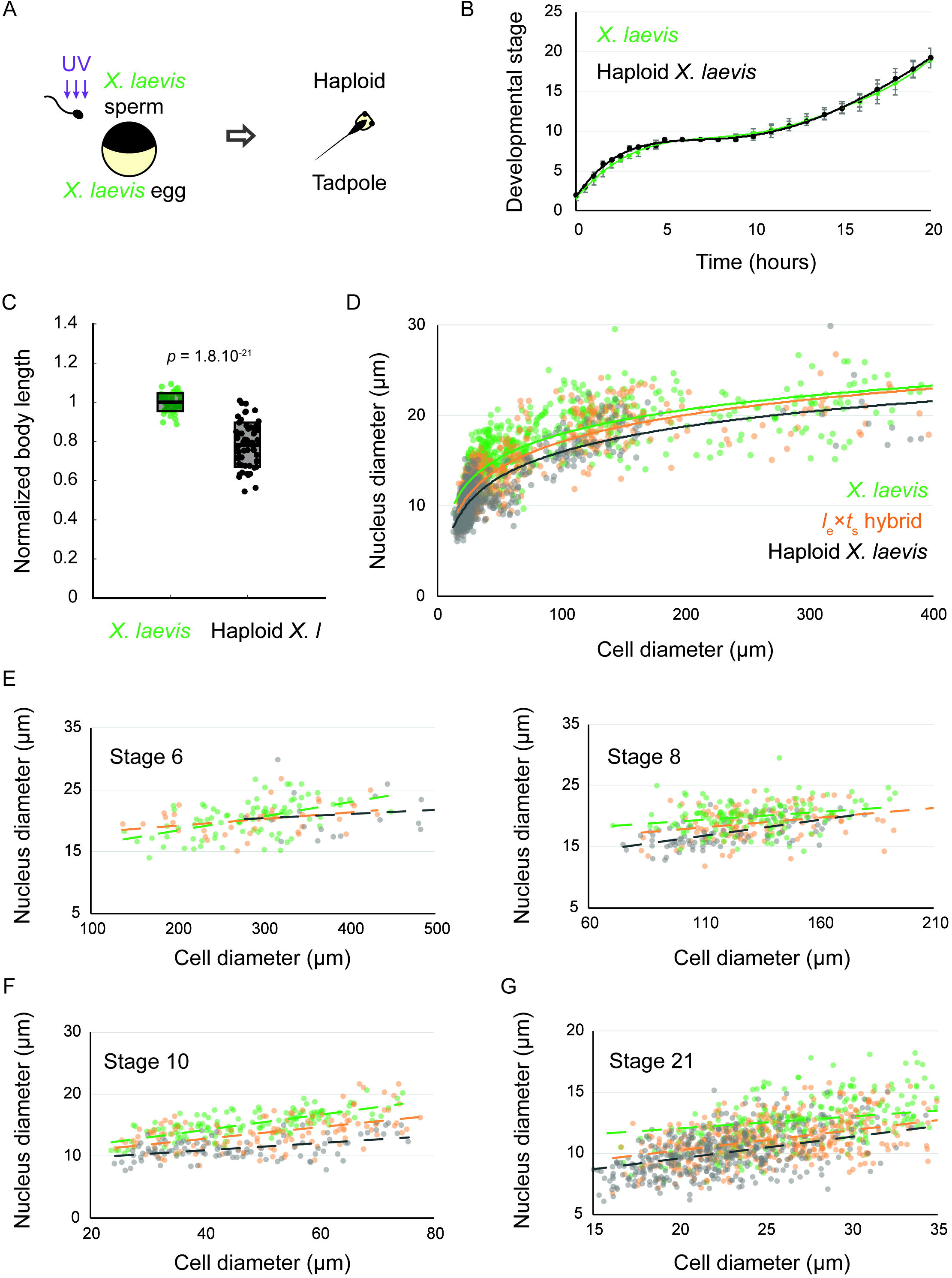
Nuclear to cell size relationships pre- and post-zygotic genome activation in *l*_e_×*t*_s_ hybrids compared to *X. laevis* diploids and haploids. **(A)** Schematic of generation of haploid *X. laevis* tadpoles via UV irradiation of sperm. **(B)** Developmental timing in *X. laevis* and haploid *X. laevis* embryos. Average is plotted for each time point. Error bars show standard deviation. **(C)** Body length of tailbud stage *X. laevis* and haploid *X. laevis*. Box plots show all individual body lengths. Thick line inside box = average length, upper and lower box boundaries = +/- SD. P-value was determined by two-tailed heteroscedastic t-test. **(D)** Nuclear diameter versus cell diameter in *X. laevis, X. laevis* haploid, and *l*_e_×*t*_s_ hybrid embryos. **(E)** Nuclear diameter versus cell diameter in *X. laevis, X. laevis* haploid, and *l*_e_×*t*_s_ hybrid embryos at developmental stages 6 and 8. **(F)** Nuclear diameter versus cell diameter in *X. laevis, X. laevis* haploid, and **l*_e_×*t*_s_* hybrid embryos at developmental stage 10. **(G)** Nuclear diameter versus cell diameter in *X. laevis, X. laevis* haploid, and *l*_e_×*t*_s_ hybrid embryos at developmental stage 21. For E-G, we ran an analysis of covariance (ANOCOVA test) to determine whether the nuclear to cell size scaling significantly depends on the embryo types. At stage 6, *p* = 0.132, at stage 8, *p* = 0.126, at stage 10, *p* = 2.558×10^−6^, and at stage 21, *p* = 1.110×10^−7^.

We then evaluated nuclear to cell size scaling relationships before and after ZGA, comparing *X. laevis, l*_e_×*t*_s_ hybrids and haploid *X. laevis* embryos (Figure 2D-G). Interestingly, no difference in the nuclear to cell size ratio was observed at early stages (6 and 8) among the 3 embryo types (Figure 2E). However, we found that, from stage 10, haploid embryos possessed reduced nuclear sizes at similar cell sizes compared to *X. laevis* (Figure 2F). Consistent with their intermediate genome size (36 > 28 > 18 chromosomes), the scaling curve of hybrids fell between that of *X. laevis* and haploids. Strikingly however, by stage 21, nuclear to cell size scaling in hybrids was more similar to that of haploids than to wild type *X. laevis* (Figure 2G). Therefore, we hypothesized that upon ZGA, gene expression of the *X. tropicalis* paternal genome, rather than bulk genome size alone, contributes to the reduced size of *l*_e_×*t*_s_ hybrids.

### Transcriptome analysis identifies 12 *X. tropicalis* transcription factors expressed in hybrids

To identify paternal *X. tropicalis* genes that could contribute to size control in hybrid embryos at ZGA, we performed RNA sequencing and transcriptome analysis of embryos at stage 9. We detected many tropicalis-derived paternally expressed genes in hybrid embryos. Differential expression analysis revealed one maternally expressed *X. laevis* gene that was significantly less abundant in the *l*_e_×*t*_s_ hybrid (Figure 3A, Supplementary Table S1), and 41 paternally expressed *X. tropicalis* genes that were significantly more abundant in *l*_e_×*t*_s_ hybrid, compared to *X. laevis* embryos (Figure 3B, Supplementary Table S1). Gene ontology (GO) term analysis of differentially expressed paternal genes revealed significant overrepresentation of the molecular function ‘DNA binding’ (GO:0003677; 4.38 fold enrichment, with a 2.90e-3 false discovery rate), and of the biological process ‘transcription, DNA-templated’ (GO:0006351; 4.65 fold enrichment, with a 3.93e-04 false discovery rate). Therefore, we conclude that transcriptional regulators with DNA binding functions are significantly enriched in paternally expressed genes in hybrid embryos. To finalize our list of candidates, we used Xenbase (James-Zorn et al., 2012; Karimi et al., 2018) to validate the transcription factor function of the candidate genes. From this, we set out to screen the following 12 transcription factors, Ers10, Hes7, Mix1, Ventx2, Foxi4, Sox3, Tgif2, Klf17, Sia2, Id3, Not and Oct25, as potential paternal scaling factors.

**Figure 3.**
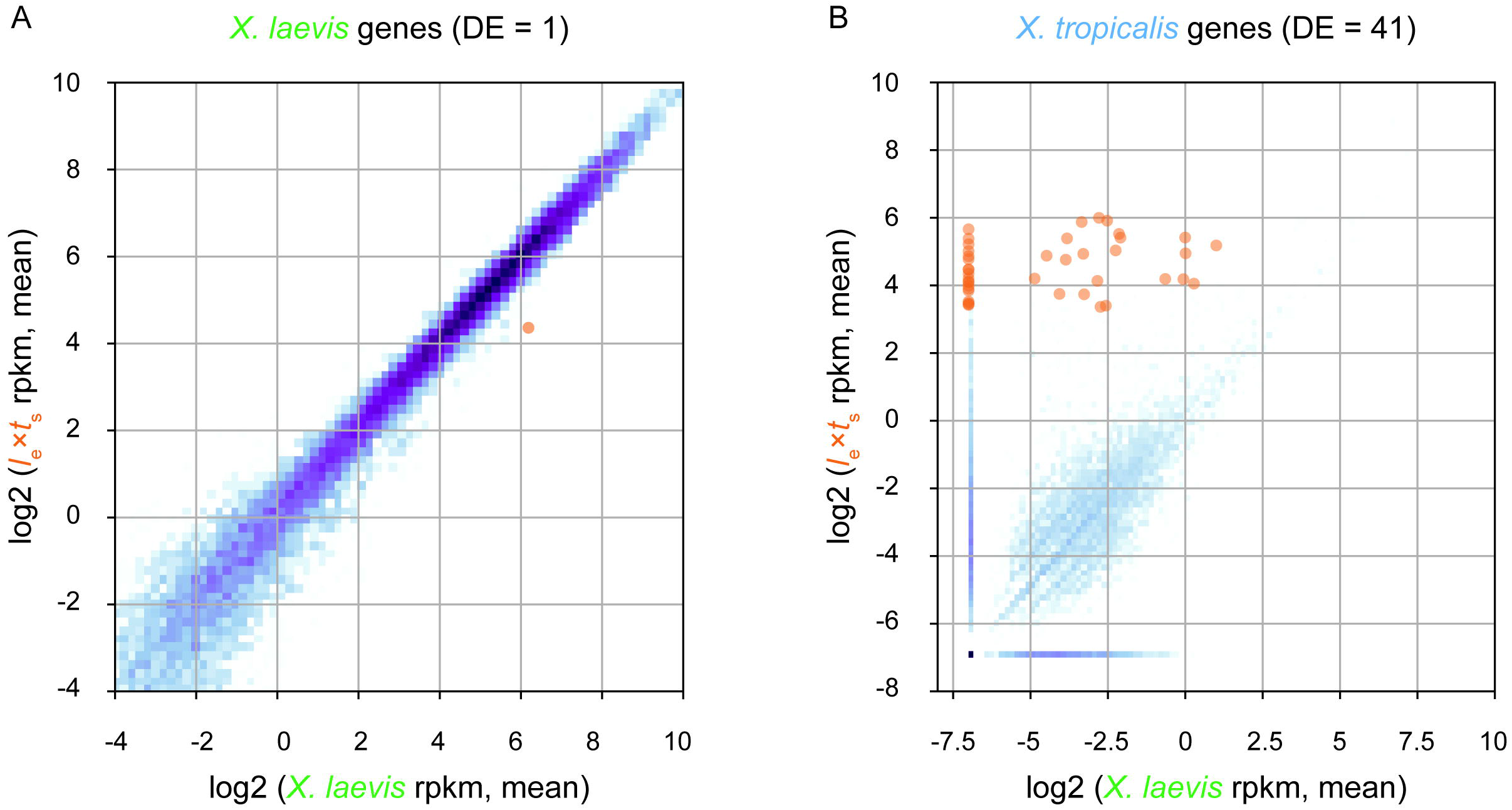
Transcriptome analysis of *l*_e_×*t*_s_ hybrid embryos at the onset of zygotic genome activation. **(A)** Differential expression analysis of *X. laevis* maternal genes in stage 9 *l*_e_×*t*_s_ hybrid vs. *X. laevis* embryos. **(B)** Differential expression analysis of *X. tropicalis* paternal genes in stage 9 *l*_e_×*t*_s_ hybrid vs. *X. laevis* embryos. For both figures, RNA-seq reads are mapped to a database of combined *X. laevis* and *X. tropicalis* transcriptomes and significantly differentially expressed genes (DE; fold-change > 2 and false discovery rate < 0.05) are marked in orange (see Material and Methods for more information).

### *X. tropicalis* transcription factors Hes7 and Ventx2 reduce body length in tailbud stage *X. laevis* embryos

To test whether identified candidate transcription factors were responsible for reducing the size of hybrid embryos, we mimicked overexpression of each transcription factor (as in the *l*_e_×*t*_s_ hybrid) by microinjecting mRNA encoding each *X. tropicalis* candidate gene into fertilized one-cell *X. laevis* embryos. Cell and nuclear size were assessed in embryos fixed for immunofluorescence around the time of ZGA (10 hours post-fertilization, ~ stage 10) and several hours post ZGA (24 hours post-fertilization, ~ stage 21). Head to tail body length was measured at late tailbud stage, 48 hours post-fertilization (Figure 4A). Two candidate transcription factors, Hes7 and Ventx2, significantly reduced overall body length (Figure 4B, C). Interestingly, the body length of embryos injected with Hes7 or Ventx2 was not significantly different from the body length in the *l*_e_×*t*_s_ hybrid (*p* = 0.64 and *p* = 0.48, respectively; two-tailed heteroscedastic t-test). It is not clear whether this effect is due to the level of overexpression, or to sequences differences between *X. laevis* and *X. tropicalis* proteins. Sequence comparisons revealed that *X. laevis* homeologs from L and S chromosomes share approximately 90% similarity, while the *X. tropicalis* Ventx2 and Hes7 from *X. tropicalis* are ~85% similar to the *X. laevis* proteins (Supplementary Figure S1A-B). The reduction in body length was however not accompanied by a change in nuclear to cell size scaling relationships as significant as that observed in the *l*_e_×*t*_s_ hybrid (Supplementary Figure S2A-B). To test whether co-expression of both genes had an additive or synergistic effect, Hes7 and Ventx2 were co-injected. This caused embryo death (27.39 ± 9.25 % lethality on average) with viable embryos more similar in size distribution to Ventx2 than to Hes7-injected embryos (Supplementary Figure S2C; *p* = 0.18 and *p* = 0.04, respectively, two-tailed heteroscedastic t-test). We also observed significant embryo death in Sia2-injected embryos (to 100% by 48 hours post-fertilization), preventing measurement at tailbud stage. Embryo death may be due to higher levels of Sia2 expression in injected embryos compared to the hybrid. However, neither nuclear nor cell size at stage 10 or 21 was altered as observed in hybrid embryos, indicating that death was not due to size scaling defects (Supplementary Figure S2D). Altogether, while the screen did not reveal factors that significantly affected cell and nuclear size, overexpression of either *X. tropicalis* Hes7 and Ventx2 resulted in a decrease in embryo size that could potentially contribute to organism size scaling in *l*_e_×*t*_s_ hybrids.

**Figure 4.**
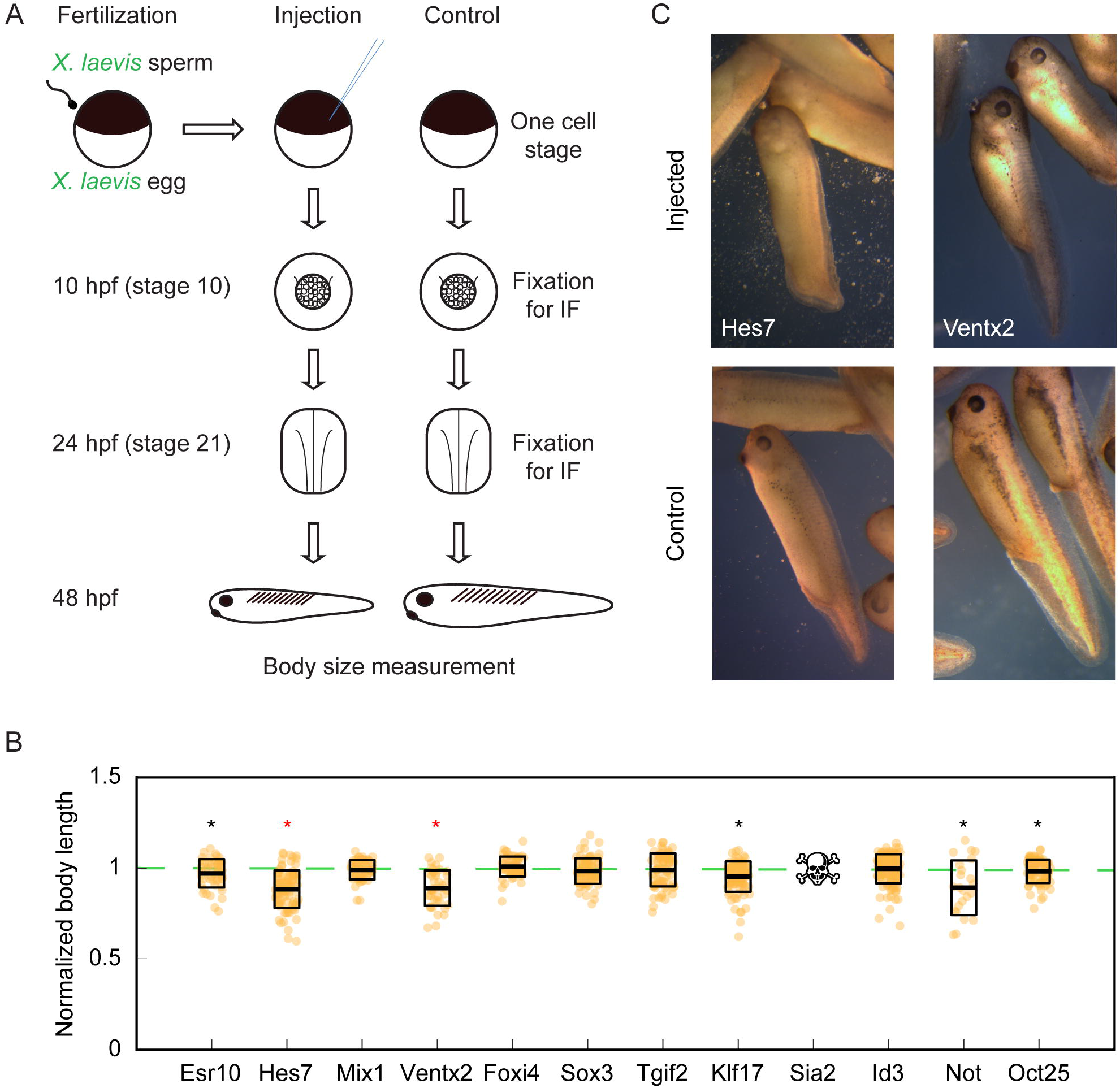
Organismal size in *X. laevis* embryos upon overexpression of candidate *X. tropicalis* transcription factors. **(A)** Workflow of candidate scaling factor screen. **(B)** Body length of tailbud stage injected *X. laevis* embryos. Thick line inside box = average length, upper and lower box boundaries = +/- SD. Stars indicated overall (results of 3 experiments pooled) significance of *p* < 0.05 (two-tailed heteroscedastic t-test). Red coloring indicates significance of each 3 individual technical replicates with *p* < 0.05 (two-tailed heteroscedastic t-test). Reduced number of measured embryos in Not is due to the fact that, overall, 67.5% of injected-embryos exogastrulated, indicating a developmental defect. **(C)** Representative images of injected *X. laevis* embryos 48 hours post-fertilization. Hes7- (left) and Ventx2-injected (right) are shown (top) with corresponding controls (bottom). Images are at identical scale.

## Discussion

Little is known about how organisms scale in size and how size scaling is coordinated at the organismal, cellular, and subcellular levels. Uniquely, between the frogs *X. laevis* and *X. tropicalis*, linear size scaling is observed at the level of the genome, subcellular structures, cell, and organism. The specific factors that influence this phenomenon are unknown.

Generating viable *l*_e_×*t*_s_ hybrids intermediate in genome size, cell size, and body size between *X. laevis* and *X. tropicalis* allowed us to examine whether size scaling in *Xenopus* results from differences in genome size alone, or whether gene expression plays a role. While genome size clearly correlates with cell and organism size in *l*_e_×*t*_s_ hybrids, other factors likely influence these parameters. Even though the genome size of the *l*_e_×*t*_s_ hybrid is closer to that of a wild type *X. laevis* embryo, the nuclear to cell size scaling curve tracked more closely with that of haploid *X. laevis* embryos. Moreover, changes in nuclear to cell size ratios in *l*_e_×*t*_s_ hybrids began at zygotic genome activation, rather than in the early cleaving embryo, which lacks transcription and growth phases. It is therefore likely that size scaling in hybrids is at least in part a consequence of *X. tropicalis* paternal gene expression rather than from reduced genome size alone.

Two transcription factors from our screen, Ventx2 and Hes7, caused a significant decrease in body length, but did not cause a change in nuclear to cell size scaling as observed in the hybrid. Reduced head-to-tail body length may be caused by different mechanisms. Ventx transcription factors have been observed to maintain pluriopotency and inhibit cell differentiation during *Xenopus* embryogenesis (Scerbo et al., 2012). Therefore, overexpression may cause a developmental delay that interferes with tissue growth. Moreover, these transcription factors may affect only specific regions of the embryo. For example, in Hes7 injected embryos, tail length is shortened, whereas head and body length remain similar to *X. laevis* controls. Hes7 is essential for regulating somite segmentation in vertebrates via oscillatory expression in presomitic mesoderm. Dampening Hes7 oscillations either by deleting or overexpressing Hes7 causes somite fusion (Bessho et al., 2003; Takashima et al., 2011) which shortens tail length in mice (Hirata et al., 2004). Interestingly, mutation of Hes7 is similarly implicated in shortening tail length in cats (Xu et al., 2016) and miniaturization of dogs (Willet et al., 2015). It is also involved in human diseases such as spondylocostal dysostosis, which causes abnormal fusion of the bones of the ribs and spine, leading to a type of dwarfism characterized by a short trunk with normal-length arms and legs (Sparrow et al., 2013).

What then precisely regulates cell and subcellular scaling in *l*_e_×*t*_s_ hybrids, and how can gene expression influence these parameters? We propose a model whereby cell size in *Xenopus* is largely dictated by genome size, but can be “fine-tuned” by differential gene expression. Such differential gene expression can also influence organism size, which may be uncoupled from cell size. Our study illustrates an example of both a unique model system and a screening approach to study biological size control and scaling. Future experiments will take advantage of the improving *Xenopus* genome assemblies to identify and screen other candidate genes, particularly those involved in other biological processes such as growth factor signaling and cell proliferation.

### Author Contributions

R.H. and R.G. designed the project. R.G. and R.A. characterized and measured nuclear and cell sizes in hybrids and haploids. T.K. performed the transcriptome analysis. K.M. and R.G. cloned the candidates and conducted the screen. K.M. measured nuclear and cell sizes in injected embryos. K.M. wrote the manuscript with inputs from R.H. and R.G.

### Funding

R.G. was initially supported by EMBO long-term fellowship ALTF 836-2013 and for most of this project by Human Frontier Science Program long-term fellowship LT 0004252014-L. K.M. was supported by the National Science Foundation Graduate Research Fellowship Program and National Institutes of Health training grant 2T32GM007232-36. R.A. was supported in part by a National Science Foundation REU Summer Fellowship in 2014. T.K. was supported by the Basic Science Research Program through the National Research Foundation of Korea funded by the Ministry of Science, ICT and Future Planning (NRF-2016R1C1B2009302), and the UNIST Research Fund (grant number 1.180063.01). R.H. was supported by NIH MIRA grant R35 GM118183 and the Flora Lamson Hewlett Chair. The confocal microscopy performed in this work was done at the UC Berkeley CRL Molecular Imaging Center, supported by National Science Foundation DBI-1041078.

## Acknowledgements

We thank members of the Heald laboratory, present and past, for support and discussions, especially C. Cadart for critical reading of the manuscript. We thank E. M. Marcotte and colleagues at the Genome Sequencing and Analysis Facility (UT Austin) for RNA-seq analysis, and J. Chapman for assistance designing primers to loci on chromosome 5 for *X. laevis* and *tropicalis*. We also thank students K. Shih, M. Fitzsimmons and S. Kawada for their dedicated assistance with experiments.

## Materials and Methods

### *Xenopus* frogs

All animal handling and procedures were performed according to the Animal Use Protocol approved by the UC Berkeley Animal Care and Use Committee. Mature *X. laevis* and *X. tropicalis* frogs were obtained from NASCO (Fort Atkinson, WI).

### Generation of viable *Xenopus l*_e_×*t*_s_ hybrid embryos

*X. laevis* females were primed with 100 IU of pregnant mare serum gonadotropin (PMSG, National Hormone and Peptide Program, Torrance, CA) at least 48 h before use and boosted with 500 IU of HCG (Human Chorionic Gonadotropin CG10, Sigma) 14-16 hours before experiments. *X. tropicalis* males were primed with 250 IU of HGC 24 hours before dissection. To obtain testes, *X. tropicalis* males were euthanized by anesthesia through immersion in double-distilled (dd)H_2_O containing 0.15% MS222 (tricaine) neutralized with 5□mM sodium bicarbonate before dissection. Testes were collected in Leibovitz L-15 media (Gibco – Thermo Fisher Scientific, Waltham, MA) supplemented with 10% Fetal Bovine Serum (FBS; Gibco), and stored at room temperature until fertilization. To prepare the sperm solution, one testis was added to 1 mL of ddH_2_O in a 1.5 mL microcentrifuge tube, and homogenized using scissors and a pestle. *X. laevis* females were squeezed gently to deposit eggs onto petri dishes coated with 1.5% agarose in 1/10X MMR. Any liquid in the petri dishes was removed and the eggs were fertilized with 1 mL of sperm solution per dish. Fertilized embryos were swirled in the solution to form a monolayer on the bottom of the petri dish and incubated for 10 min with the dish slanted to ensure submersion of eggs. Dishes were then flooded with 1/10X MMR, swirled and incubated for 10 min. To remove egg jelly coats, the 1/10X MMR was completely exchanged for freshly prepared Dejellying Solution (2% L-cysteine in ddH_2_O-NaOH, pH 7.8). After dejellying, eggs were washed extensively (>4X) with 1/10X MMR before incubation at 23°C. At Nieuwkoop and Faber stage 2-3, fertilized embryos were sorted and placed in fresh 1/10X MMR in new petri dishes coated with 1.5% agarose in 1/10X MMR.

### Confirmation of presence of both *X. laevis* and *X. tropicalis* genomes in *l*_e_×*t*_s_ hybrids

Genomic DNA was extracted from *l*_e_×*t*_s_ hybrid embryos by incubating overnight in lysis buffer (50 mM Tris-HCl, 5 mM EDTA, 100 mM NaCl, 0.5% SDS) containing 250 μg/mL Proteinase K (Roche, Basel, Switzerland). DNA was isolated using Phenol-Chloroform extraction and ethanol precipitation. The genomic DNA was used as a PCR template for a single pair of primers that amplify a specific locus that differs ~100bp in size between all 3 (sub)genomes. In *X. tropicalis*, the locus is on chromosome 5 and PCR product size is 510 bp. In *X. laevis*, one locus is on chromosome 5L for which PCR product size is 408 bp and another one is on chromosome 5S for which PCR product size is 305 bp. The sequences of the primer pair are fwd GTACTCTTCCCCAGCTTGCTG and rev GCCTGTATGGCTCCTAGGTTTTC.

### Generation of wild type *X. laevis* embryos for microinjection

Ovulations, euthanasias, dissections, and fertilizations were carried out as described above for *l*_e_×*t*_s_ hybrids above, with the following modifications: *X. laevis* males were primed by injecting 500 IU of HCG 24 hours before dissection. Dissected testes were collected in 1X Modified Ringer (MR) (100 mM NaCl, 1.8 mM KCl, 1 mM MgCl2, 5 mM HEPES-NaOH pH 7.6 in ddH2O), and stored at room temp for short periods, or at 4°C for up to 5 days. To make sperm solution, 1/3-1/2 of a testis was added to 1 mL of ddH_2_O in a 1.5 mL microcentrifuge tube.

### Generation of haploid *X. laevis* embryos

Euthanasia of males and dissection/collection of testes proceeded as described for *X. laevis* males above. 1/3-1/2 of a testis was added to 1.1 mL of ddH_2_O in a 1.5 mL microcentrifuge tube and homogenized with scissors and a pestle. The tube was briefly centrifuged using a benchtop microcentrifuge for several seconds to pellet large pieces of tissue. One mL of supernatant was removed, avoiding pieces of tissue, and transferred to a non-coated glass petri dish. The open dish was placed into a UV-Crosslinker (Stratalinker, Stratagene) and the sperm solution irradiated twice using 30,000 microjoules. The solution was swirled between the two irradiations. The irradiated sperm solution was then retrieved and used for fertilization by depositing at least 0.5 mL solution on top of freshly squeezed *X. laevis* eggs in a petri dish coated with 1.5% agarose in 1/10x MMR. Fertilization, dejelly, and embryo storage then proceeded as described for *l*_e_×*t*_s_ hybrid embryos above.

### Embryo video imaging

Imaging dishes were prepared using a homemade PDMS mold designed to print a pattern of 1 mm large wells in agarose that allowed us to image 4 embryos simultaneously within the 3×4 mm camera field of view for each type of embryo. Embryos were imaged from stage 2. *X. laevis* and *l*_e_×*t*_s_ hybrid or haploid videos were taken simultaneously using two AmScope MD200 USB cameras, (AmScope, Irvine, CA) each mounted on an AmScope SE305R stereoscope. Time lapse movies were acquired at a frequency of 1 frame every 10 s for 20 h and saved as Motion JPEG using a MATLAB (The MathWorks, Inc., Natick, MA) script. Movie post-processing (cropping, concatenation, resizing, addition of scale bar) was done using MATLAB and Fiji (Schindelin et al., 2012). All MATLAB scripts written for this study are available upon request. Two of the scripts used here were obtained through the MATLAB Central File Exchange: “videoMultiCrop” and “concatVideo2D” by Nikolay S.

### Imaging and measurement of tailbud, tadpole and frog body size

Tailbud stage embryos were placed in an ice-cold agarose-coated imaging chamber and imaged at 12x magnification using a Wild Heerbrugg M7A StereoZoom microscope coupled to a Leica MC170HD camera and Leica LAS X software. Tadpoles were imaged by placing in a petri dish filled with a limited amount of water to prevent depth-biased measurements. Images were taken with an iphone camera, including a ruler in the field of view. Tadpole measurements were stopped when the tail began to recede at the end of metamorphosis. Froglets were placed in a transparent-bottom container placed on a ruler and fill with a minimal amount of water, and imaged with an iphone camera. Images were analyzed and length measured head to tail for tadpoles, or head to cloaca for froglets. Length measurements were done using the line tool in Fiji.

### Erythrocyte preparation and measurements

A small drop of blood was collected from the frog foot with a sterile needle, and the drop was smeared on a slide. The smear was then fixed with methanol and stained with Giemsa stain (Sigma GS). Cell were imaged in brightfield using micromanager software (Edelstein et al., 2014) with an Olympus BX51 microscope equipped with an ORCA-II camera (Hamamatsu Photonics, Hamamatsu city, Japan).

### RNA isolation and sequencing

To isolate RNA, embryos at stage 9 were homogenized mechanically in TRIzol^®^ (Thermo Fisher Scientific, Waltham, MA) using up to a 30-gauge needle and processed according to manufacturer instructions. After resuspension in nuclease-free H2O, RNAs were cleaned using a RNeasy kit (Qiagen Inc.) according to manufacturer instructions. Libraries were prepared using manufacturer’s non-standard specific RNA-seq library protocol with poly-A capturing mRNA enrichment method (Illumina, CA, USA). The paired-end 2 × 100 bp reads were generated by the Genome Sequencing and Analysis Facility (GSAF) at the University of Texas at Austin using Illumina HiSeq 2000. Transcriptome data generated in this study are available from NCBI Gene Expression Omnibus (Series record GSE118382).

### Gene Expression Analysis

We mapped RNA-seq reads to the database of combined *X. laevis* and *X. tropicalis* transcriptome (available at http://genome.taejoonlab.org/pub/xenopus/annotation/; WorldCup_201407 version), using Bowtie1 (version 1.0). To prevent misalignment to other species, we applied stringent criteria, allowing no mismatches (-v 0), and ignoring a read mapped more than one target (-m 1). We estimated relative transcript abundance with ‘transcripts per million reads (TPM)’ calculated by RSEM (version 1.2.19), and differential expression analysis was conducted using edgeR (version 3.36.1), with greater than two-fold changes and false discovery rate (FDR) less than 0.05 cutoff to determine the significance.

### Gene Ontology Analysis

We conducted Gene Ontology analysis with Panther DB (version 13.1). For statistical analysis for overrepresented terms, we used Fischer’s exact test and FDR adjustment, and applied FDR less than 0.05 as a significance cutoff. To validate our list of candidates, we searched Xenbase (http://www.xenbase.org/) using the gene name as the query.

### Cloning and mRNA synthesis of candidate transcription factors

Total RNA was isolated from *X. tropicalis* embryos as described above in “RNA isolation and sequencing”, and cDNA was synthesized from RNA using the SuperScript III First Strand Synthesis system (Invitrogen-Thermo Fisher Scientific, Waltham, MA) according to manufacturer instructions. Transcription factor sequences were then PCR-amplified from the cDNA using the following primer sequences (all are written 5’-3’) concatenated with ~30 bp plasmid-homologous sequences: Esr10, fwd ATGGCTCCTTACAGCGCTAC, rev TTCTCTGGAGACCCTGGAAC; Sox3, fwd ATGTATAGCATGTTGGACAC, rev CTGTACCGCTCACTCACATA; Foxi4.2, fwd ATGAACCCAGTCCAGCAACC, rev CTTTGTACCAGGGAAGGTAC; Hes7.1, fwd AT GA AGGGAGC GAGT GA AGT, rev AGACCTGGAGACCTTGGGTA; Mix1, fwd ATGGACTCATTCAGCCAACA, rev TCTGTGTGCTCCTCCACCTT; Tgif2, fwd ATGATGAATTCGACTTTTGA, rev TCACGACAAGCACCCCCAAT; Ventx2.1, fwd ATGAACACAAGGACTACTAC, rev TTGGGCAGCCTCTGGCCTAC; Klf17, fwd ATGAGTGTGGCTTTCTCAAC, rev CATGTGTCTCTTCATGTGCAG; Not, fwd ATGTTACACAGCCCTGTCTTTC, rev CAGTTCAACATCCACATCATC; Oct25 fwd ATGTACAGCCAACAGCCCTTC, rev ACCAATATGGCCGCCCATGG; Sia2 fwd ATGACTTGTGACTCTGAGCTTG, rev GCCCCACATATCCGGATATTG; Id3 fwd ATGAAAGCCATCAGCCCAGTG, rev GTGGCAGACACTGGCGTCCC. These amplified sequences were then subcloned using Gibson assembly (New England Biolabs, Ipswich, MA) into a PCS2 expression vector obtained at the 2013 Advanced Imaging in *Xenopus* Workshop from the Wallingford lab (UT Austin, USA). mRNAs were synthetized from these expression constructs using mMessage mMachine SP6 Transcription Kit (Ambion – Thermo Fisher Scientific, Waltham, MA) following the manufacturer protocol. The mRNAs were then purified using Phenol-Chloroform extraction, resuspended in ddH_2_O, aliquoted and stored at −80°C.

### Microinjection of candidate transcription factors into *Xenopus* embryos

Stage 1 (one-cell) embryos about 30 minutes post-fertilization were transferred into a mesh-bottom dish containing 1/9X MMR 3% Ficoll for microinjection. Injections were done using a Picospritzer III microinjection system (Parker, Hollis, NH) equipped with a MM-3 micromanipulator (Narishige, Amityville, NY). To mimic overexpression of each transcription factor, each embryo was injected with 750 picograms of mRNA, a dose we determined was large enough to see phenotypes, but was not associated with embryo toxicity. Injected embryos were transferred to a new dish coated with 1.5% agarose in 1/10x MMR, and incubated at 23°C in 1/9X MMR 3% Ficoll for at least 6 hours. The embryos were then transferred to fresh 1/10x MMR in a new agarose-coated dish, and incubated at 23°C with buffer changes into fresh 1/10x MMR several times daily until ready for fixation or imaging.

### Embryo whole mount immunofluorescence

Embryos at the desired developmental stage were fixed for one hour using MAD fixative (2 parts methanol [Thermo Fisher Scientific, Waltham, MA], 2 parts acetone [Thermo Fisher Scientific, Waltham, MA]), 1 part DMSO [Sigma]). After fixation, embryos were dehydrated in methanol and stored at −20°C. Embryos were then processed as previously described (Lee et al., 2008) with modifications. Following gradual rehydration in 0.5X SSC (1X SSC: 150 mM NaCl, 15 mM Na citrate, pH 7.0), embryos were bleached with 1-2% H_2_O_2_ (Thermo Fisher Scientific, Waltham, MA) in 0.5X SSC containing 5% formamide (Sigma) for 2-3 h under light, then washed in PBT (137 mM NaCl, 2.7 mM KCl, 10 mM Na_2_HPO_4_, 0.1% Triton X-100 [Thermo Fisher Scientific, Waltham, MA]) and 2 mg/mL bovine serum albumin (BSA). Embryos were blocked in PBT supplemented with 10% goat serum (Gibco – Thermo Fisher Scientific, Waltham, MA) and 5% DMSO for 1-3 h and incubated overnight at 4°C in PBT supplemented with 10% goat serum and primary antibodies. The following antibodies were used to label tubulin and DNA, respectively: 1:500 mouse anti-beta tubulin (E7; Developmental Studies Hybridoma Bank, Iowa City, IA), and 1:500 rabbit anti-histone H3 (ab1791; Abcam, Cambridge, MA). Embryos were then washed 4× 2 h in PBT and incubated overnight in PBT supplemented with 1:500 goat anti-mouse or goat anti-rabbit secondary antibodies coupled either to Alexa Fluor 488 or 568 (Invitrogen – Thermo Fisher Scientific, Waltham, MA). Embryos were then washed 4× 2 h in PBT and gradually dehydrated in methanol. Embryos were cleared in Murray’s clearing medium (2 parts of Benzyl Benzoate, 1 part of Benzyl Alcohol).

### Confocal imaging and measurement of embryos, cells and nuclei after whole mount immunofluorescence

Embryos were placed in a chamber made using a flat nylon washer (Grainger, Lake Forest, IL) attached with nail polish (Sally Hansen, New York, NY) to a slide, filled with Murray’s clearing medium, and covered by a coverslip (Beckman coulter, Brea, CA) for confocal microscopy. Confocal microscopy was performed on a Zeiss LSM 780 NLO AxioExaminer running the Zeiss Zen Software. Embryos were imaged using a Plan-Apochromat 20x/1.0 water objective and laser power of 12%, on multiple 1024×1024 pixel plans spaced 0.68 μm apart in Z.

Nuclear area was measured in Fiji using the ellipse tool. From this, we calculated the diameter of a circle of the same area, a value that we could directly compare the cell size determined through the measurement of the cell diameter at the nucleus central plane. To test whether the nuclear to cell size scaling significantly depends on the embryo types or whether embryos were microinjected or not, we ran an analysis of covariance (ANOCOVA test) using the ‘aoctool’ in MATLAB with a ‘separate lines’ model.

### Protein sequence alignments

Multiple sequence alignments were performed using Clustal Omega (default parameters). Sequence identities and similarities were determined by pairwise alignments using EMBOSS Needle (default parameters).

## References

Bessho, Y., Hirata, H., Masamizu, Y., and Kageyama, R. (2003). Periodic repression by the bHLH factor Hes7 is an essential mechanism for the somite segmentation clock. Genes Dev. 17, 1451–1456. doi:10.1101/gad.1092303.

Björklund, M., Taipale, M., Varjosalo, M., Saharinen, J., Lahdenperä, J., and Taipale, J. (2006). Identification of pathways regulating cell size and cell-cycle progression by RNAi. Nature 439, 1009–13. doi:10.1038/nature04469.

Brown, K. S., Blower, M. D., Maresca, T. J., Grammer, T. C., Harland, R. M., and Heald, R. (2007). Xenopus tropicalis egg extracts provide insight into scaling of the mitotic spindle. J. Cell Biol. 176, 765–770. doi:10.1083/jcb.200610043.

Bürki, E. (1985). The expression of creatine kinase isozymes in Xenopus tropicalis, Xenopus laevis laevis, and their viable hybrid. Biochem. Genet. 23, 73–88.

Cavalier-Smith, T. (2005). Economy, speed and size matter: Evolutionary forces driving nuclear genome miniaturization and expansion. in Annals of Botany, 147–175. doi:10.1093/aob/mci010.

Edelstein, A. D., Tsuchida, M. a, Amodaj, N., Pinkard, H., Vale, R. D., and Stuurman, N. (2014). Advanced methods of microscope control using μManager software. J. Biol. Methods 1, 10. doi:10.14440/jbm.2014.36.

Elurbe, D. M., Paranjpe, S. S., Georgiou, G., van Kruijsbergen, I., Bogdanovic, O., Gibeaux, R., et al. (2017). Regulatory remodeling in the allo-tetraploid frog Xenopus laevis. Genome Biol. 18, 198. doi:10.1186/s13059-017-1335-7.

Frawley, L. E., and Orr-Weaver, T. L. (2015). Polyploidy. Curr. Biol. 25, R353–R358. doi:10.1016/j.cub.2015.03.037.

Galitski, T., Saldanha, A. J., Styles, C. A., Lander, E. S., and Fink, G. R. (1999). Ploidy regulation of gene expression. Science (80-.). 285, 251–4. doi:10.1126/science.285.5425.251.

Gibeaux, R., Acker, R., Kitaoka, M., Georgiou, G., Van Kruijsbergen, I., Ford, B., et al. (2018). Paternal chromosome loss and metabolic crisis contribute to hybrid inviability in Xenopus. Nature 553, 337–341. doi:10.1038/nature25188.

Good, M. C., Vahey, M. D., Skandarajah, A., Fletcher, D. A., and Heald, R. (2013). Cytoplasmic volume modulates spindle size during embryogenesis. Science (80-.). 342, 856–860. doi:10.1126/science.1243147.

Gregory, T. R. (2001). Coincidence, coevolution, or causation? DNA content, cell size, and the C-value enigma. Biol. Rev. Camb. Philos. Soc. 76, 65–101. doi:10.1111/j.1469-185X.2000.tb00059.x.

Hamilton, L. (1957). Androgenic haploids of a toad, Xenopus laevis. Nature 179, 159. doi:10.1038/179159a0.

Hamilton, L. (1963). An experimental analysis of the development of the haploid syndrome in embryos of Xenopus laevis. J. Embryol. Exp. Morphol. 11, 267–78.

Hirata, H., Bessho, Y., Kokubu, H., Masamizu, Y., Yamada, S., Lewis, J., et al. (2004). Instability of Hes7 protein is crucial for the somite segmentation clock. Nat. Genet. 36, 750–754. doi:10.1038/ng1372.

James-Zorn, C., Ponferrada, V. G., Jarabek, C. J., Burns, K. a, Segerdell, E. J., Lee, J., et al. (2012). Xenbase: expansion and updates of the Xenopus model organism database. Nucleic Acids Res., 1–6. doi:10.1093/nar/gks1025.

Karimi, K., Fortriede, J. D., Lotay, V. S., Burns, K. A., Wang, D. Z., Fisher, M. E., et al. (2018). Xenbase: a genomic, epigenomic and transcriptomic model organism database. Nucleic Acids Res. 46, D861–D868. doi:10.1093/nar/gkx936.

Kitaoka, M., Heald, R., and Gibeaux, R. (2018). Spindle assembly in egg extracts of the Marsabit clawed frog, Xenopus borealis. Cytoskeleton. doi:10.1002/cm.21444.

Lee, C., Kieserman, E., Gray, R. S., Park, T. J., and Wallingford, J. (2008). Whole-mount fluorescence immunocytochemistry on Xenopus embryos. CSH Protoc. 2008, pdb.prot4957. doi:10.1101/pdb.prot4957.

Lee, H. O., Davidson, J. M., and Duronio, R. J. (2009). Endoreplication: Polyploidy with purpose. Genes Dev. 23, 2461–2477. doi:10.1101/gad.1829209.

Levy, D. L., and Heald, R. (2010). Nuclear Size Is Regulated by Importin α and Ntf2 in Xenopus. Cell 143, 288–298. doi:10.1016/j.cell.2010.09.012.

Lindsay, L. L., Peavy, T. R., Lejano, R. S., and Hedrick, J. L. (2003). Cross-fertilization and structural comparison of egg extracellular matrix glycoproteins from Xenopus laevis and Xenopus tropicalis. Comp. Biochem. Physiol. A. Mol. Integr. Physiol. 136, 343–52. doi:10.1016/S1095-6433(03)00169-7.

Loughlin, R., Wilbur, J. D., McNally, F. J., Nedelec, F. J., and Heald, R. (2011). Katanin contributes to interspecies spindle length scaling in xenopus. Cell 147, 1397–1407. doi:10.1016/j.cell.2011.11.014.

Marguerat, S., and Bähler, J. (2012). Coordinating genome expression with cell size. Trends Genet. 28, 560–565. doi:10.1016/j.tig.2012.07.003.

Marguerat, S., Schmidt, A., Codlin, S., Chen, W., Aebersold, R., and Bähler, J. (2012). Quantitative analysis of fission yeast transcriptomes and proteomes in proliferating and quiescent cells. Cell 151, 671–683. doi:10.1016/j.cell.2012.09.019.

Mirsky, A. E. (1951). THE DESOXYRIBONUCLEIC ACID CONTENT OF ANIMAL CELLS AND ITS EVOLUTIONARY SIGNIFICANCE. J. Gen. Physiol. 34, 451–462. doi:10.1085/jgp.34.4.451.

Narbonne, P., Simpson, D. E., and Gurdon, J. B. (2011). Deficient induction response in a Xenopus nucleocytoplasmic hybrid. PLoS Biol. 9. doi:10.1371/journal.pbio.1001197.

Neumann, F. R., and Nurse, P. (2007). Nuclear size control in fission yeast. J. Cell Biol. 179, 593–600. doi:10.1083/jcb.200708054.

Nieuwkoop, P. D., and Faber, J. (1994). Normal table of Xenopus laevis (Daudin). Garland Publishing.

Scerbo, P., Girardot, F., Vivien, C., Markov, G. V., Luxardi, G., Demeneix, B., et al. (2012). Ventx factors function as Nanog-like guardians of developmental potential in Xenopus. PLoS One 7. doi:10.1371/journal.pone.0036855.

Schindelin, J., Arganda-Carreras, I., Frise, E., Kaynig, V., Longair, M., Pietzsch, T., et al. (2012). Fiji: An open-source platform for biological-image analysis. Nat. Methods 9, 676–682. doi:10.1038/nmeth.2019.

Sparrow, D. B., Faqeih, E. A., Sallout, B., Alswaid, A., Ababneh, F., Al-Sayed, M., et al. (2013). Mutation of HES7 in a large extended family with spondylocostal dysostosis and dextrocardia with situs inversus. Am. J. Med. Genet. Part A 161, 2244–2249. doi:10.1002/ajmg.a.36073.

Taft, R. J., Pheasant, M., and Mattick, J. S. (2007). The relationship between non-protein-coding DNA and eukaryotic complexity. BioEssays 29, 288–299. doi:10.1002/bies.20544.

Takashima, Y., Ohtsuka, T., Gonzalez, A., Miyachi, H., and Kageyama, R. (2011). Intronic delay is essential for oscillatory expression in the segmentation clock. Proc. Natl. Acad. Sci. 108, 3300–3305. doi:10.1073/pnas.1014418108.

Wilbur, J. D., and Heald, R. (2013). Mitotic spindle scaling during Xenopus development by kif2a and importin α. Elife 2, e00290. doi:10.7554/eLife.00290.

Willet, C. E., Makara, M., Reppas, G., Tsoukalas, G., Malik, R., Haase, B., et al. (2015). Canine disorder mirrors human disease: Exonic deletion in HES7 causes autosomal recessive spondylocostal dysostosis in miniature schnauzer dogs. PLoS One 10. doi:10.1371/journal.pone.0117055.

Xu, X., Sun, X., Hu, X. S., Zhuang, Y., Liu, Y. C., Meng, H., et al. (2016). Whole Genome Sequencing Identifies a Missense Mutation in HES7 Associated with Short Tails in Asian Domestic Cats. Sci. Rep. 6. doi:10.1038/srep31583.

